# Learning Mappings from Cryo-EM Images to Atomic Coordinates via Latent Representations

**DOI:** 10.64898/2026.02.16.706167

**Authors:** Eya Abid, Slavica Jonic

## Abstract

Single-particle cryo-electron microscopy (cryo-EM) aims to determine three-dimensional (3D) structures of biomolecular complexes from noisy two-dimensional (2D) projection images acquired at unknown orientations. The presence of pose uncertainty and continuous conformational heterogeneity makes high-resolution reconstruction challenging. Here, we investigate, in a controlled synthetic setting, whether supervised learning can map noisy cryo-EM single-particle images to atomic coordinates without pose recovery or 2D projection calculations. We propose a convolutional auto-encoder to compress particle images into their corresponding latent representations, followed by a regression network to predict 3D atomic coordinates from these image latents. We show the performance of this approach using synthetic datasets of pairs of particle images and conformational models of adenylate kinase and nucleosome core particles, generated using a realistic cryo-EM forward model based on Normal Mode Analysis for simulating dynamics. Inference yielded mean RMSDs of 2.11 Å for all-atom models of adenylate kinase (1,656 atoms) and 0.80 Å for the coarse-grain models of nucleosome (1,041 Cα-P atoms). These results indicate that compact image latents preserve pose and conformation related information sufficiently well to support atomic coordinate regression. This provides a quantitative proof-of-principle for coupling image and structure spaces toward fast estimation of conformational variability in cryo-EM.

## I. Introduction

Cryo-electron microscopy (cryo-EM) enables structural characterization of biomolecular assemblies in near-native conditions by reconstructing 3D density maps from collections of noisy 2D projection images [1, 2]. In typical experiments, each particle is imaged at an unknown orientation, and images are corrupted by noise and contrast transfer function (CTF) effects. Many complexes also populate continuous conformational landscapes rather than discrete states [3, 4]. Continuous conformational trajectories have been observed in experimental cryo-EM datasets, for example in studies of ribosome dynamics [5]. Thus, reconstruction requires addressing a coupled problem of pose estimation and conformational variability under severe noise.

Conventional pipelines alternate between pose estimation, classification, averaging, and refinement. While effective for rigid complexes, their performance degrades as conformational variability becomes continuous or when the signal-to-noise ratio (SNR) is low. Alternative approaches have explored the integration of physical priors and machine-learning models to characterize continuous conformational variability from cryo-EM images. They include flexible fitting of particle images based on normal modes or classical molecular dynamics (MD) simulations [6, 7], which result in reduced-mechanics conformational landscapes [6] or full-MD conformational landscapes [7]. Also, approaches based on neural-network latent representations and statistical machine learning have been proposed [5, 8-11], but they usually result in density-based conformational landscapes [9, 11] and rarely in atomistic conformational landscapes [10].

Among the most popular methods are CryoDRGN [9] and 3DFlex [11]. CryoDRGN leverages deep generative modeling to recover heterogeneous 3D density distributions from particle images, whereas 3DFlex estimates continuous structural variability by deforming a consensus 3D density map. Deep learning strategies have also been proposed to accelerate physically grounded pipelines such as HEMNMA, which uses normal modes to analyze particle images [6, 12]. For instance, DeepHEMNMA accelerates HEMNMA more than 40 times using a ResNet-based predictor [10]. DeepHEMNMA predicts a few conformational parameters, which are amplitudes of a few selected low-frequency normal modes, along with the pose parameters (three Euler angles and two in-plane shifts), for each particle image.

In contrast to DeepHEMNMA, the supervised neural network approach that is presented in this article aims to predict many conformational parameters, which are Cartesian atomic coordinates for each particle, with the particle poses integrated into the coordinates. In the present work, we evaluate, in a controlled setting with synthetic data, whether an image latent representation, extracted by a convolutional autoencoder, can support a direct regression of atomic coordinates, without pose estimation or comparing 2D projections of predicted atomic models with given images.

To address this question, we conduct a fully controlled study in which ground-truth atomic coordinates are known. We assess the accuracy of atomic model reconstruction with our new neural-based approach using synthetic training and test datasets, which consist of pairs of synthesized atomic coordinates and corresponding synthetic images, for two complexes of different sizes, architectures, and conformational dynamics (adenylate kinase and nucleosome core particle).

In the case of supervised training with experimental cryo-EM images, the per-particle ground-truth atomic coordinates would be unavailable, but they could be estimated using MDSPACE [7]. MDSPACE derives per-particle atomic models by analyzing particle images using flexible fitting based on classical MD simulations (based on Newton’s equations of motion). High computational cost of MD simulations makes the analysis of large datasets with MDSPACE slow. A combination of MDSPACE with the neural approach described here will be described in a separate article to show how the neural models trained with atomic models derived by MDSPACE from small datasets could be used for fast prediction of atomic coordinates on large datasets that are unamenable to MDSPACE.

The new neural approach is presented in section II. Section III describes the data synthesis procedure. The evaluation of the performance of this novel approach using synthetic data is presented in section IV, which is followed by a discussion and a conclusion (sections V and VI).

## II. Methods

Our framework consists of a combination of two networks: an image autoencoder that produces a compact latent representation (step 1, **Fig. 1**) and a regressor that maps the latent to atomic coordinates (step 2, **Fig. 2**).

**Fig. 1.**
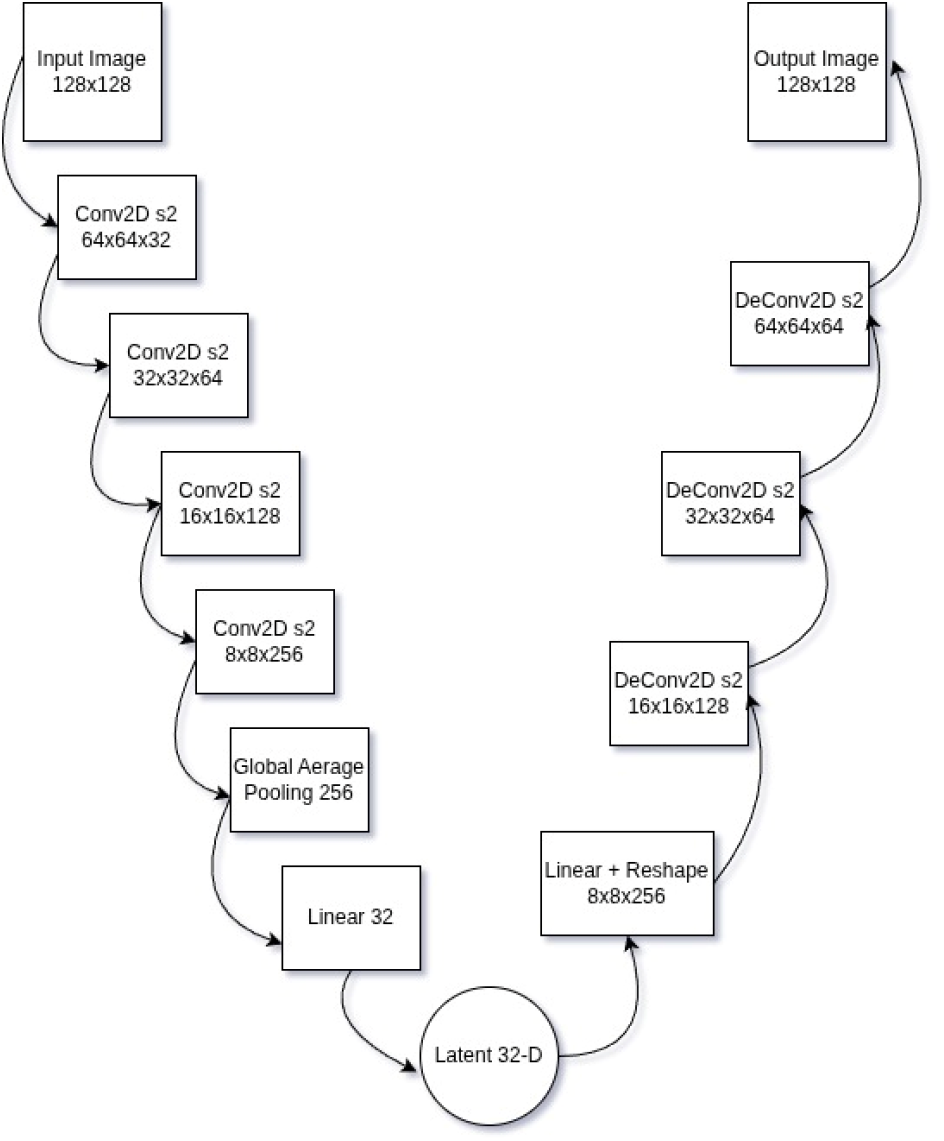
Autoencoder architecture. The encoder compresses each 128×128 pixels image into a 32-dimensional latent via four convolutional down-sampling blocks and global average pooling. The decoder reconstructs the image through a symmetric stack of transposed convolutions.

**Fig. 2.**
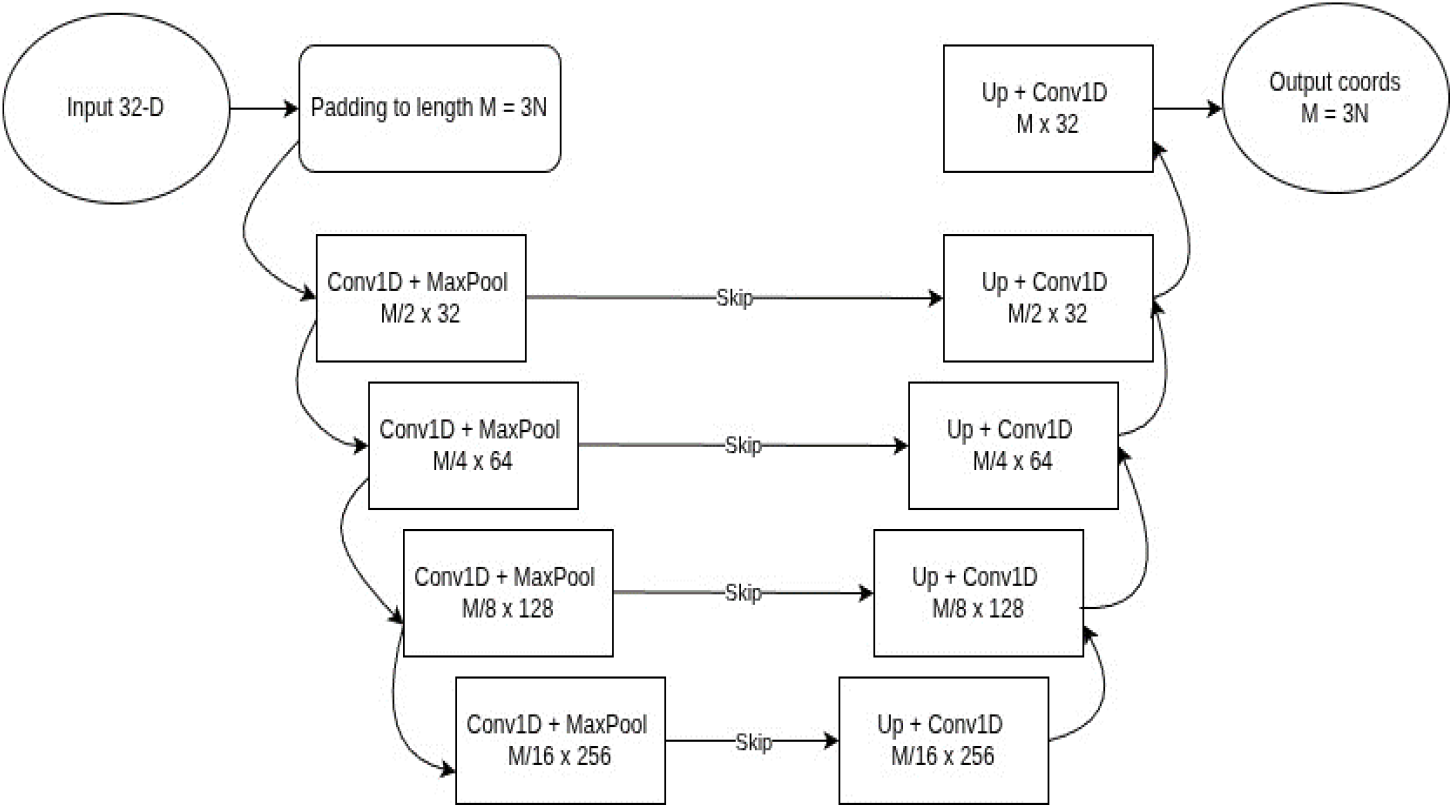
Image-to-Atoms regression network. A U-Net-inspired architecture reduces a zero-padded image latent vector of length *M* × 1 (*M = 3N* for *N* atoms) into a low-dimensional representation (length *M/16*) and expands it to *M* coordinates. The image latent is first zero-padded from length 32 × 1 (32-D) to length *M* × 1 (*M* >> 32). Skip connections improve fine-scale accuracy.

### A. Step 1: Image Autoencoder

In this article, we assume that particle images have a size of 128×128 pixels. The input to the encoder are single-channel images with pixel intensities in the range [0,1], normalized using per-image min-max normalization. The encoder applies four stride-2 3×3 convolutions, halving image dimensions at each step while increasing channel depth as follows: 128×128×1 → 64×64×32 → 32×32×64 → 16×16×128 → 8×8×256 (the first two dimensions stand for image dimensions and the third dimension is the number of channels). Global average pooling collapses the 8×8×256 tensor to a vector of 256 dimensions (256-D), then a linear layer maps it to a latent of 32 dimensions (32-D) named *z*. The decoder mirrors this process: a linear layer reshapes *z* into 8×8×256, then four stride-2 transposed convolutions restore spatial resolution and image dimensions to 128×128×1. The model is trained with Mean Squared Error (MSE) between the input image and reconstructed image as the loss, Adam optimizer (initial learning rate 0.001 with cosine decay), batch sizes 4, and 200-300 epochs. The convolutional autoencoder architecture is illustrated in **Fig. 1**. The latents of all training images are then used for regression at step 2.

### B. Step 2: Image-to-Atoms Regressor

The image-to-atom regressor is a 1D U-Net that maps each image latent *z* of length 32 (32-D) to the corresponding 3*N* (=*M*) Cartesian atomic coordinates, for a system with *N* atoms (*M*>> 32). The image latent is first expanded from length 32 to length *M*, by duplicating the latent three times (for the *x, y*, and *z* coordinates) and padding the end of each of them with zeros, followed by combining the three padded copies in a single vector of length *M*. The encoder consists of four blocks of 1D convolutions and max pooling that progressively down-sample the padded-version of the image latent (length *M*) while increasing the number of channels: *M* × 1 → *M*/*2* × 32 → *M*/*4* × 64→ *M*/*8* × 128 → *M*/*16* × 256 (the first dimension stand for signal dimension and the second dimension is the number of channels). The decoder mirrors this, with blocks consisting of up-sampling and 1D convolutions, back to length *M*, with skip connections between matching scales. Each block uses layer normalization, Gaussian Error Linear Unit (GELU) activation, and dropout (0.1-0.2). GELU is an activation function that smoothly weights inputs by their probability under a Gaussian distribution, behaving like a softer, probabilistic version of ReLU. Skip connections preserve low and mid-scale information across the expansion and contraction paths.

The output layer is linear and directly regresses coordinates without alignment. Let ***r*_*i*_** in *R*^*3*^ denote the vector of ground-truth coordinate of atom *i* and ***p*_*i*_** the vector of predicted coordinates. The training loss for the coordinate regressor is Root Mean Square Deviation (RMSD), which is a commonly used metric to evaluate the distance between atomic models in structural biology:

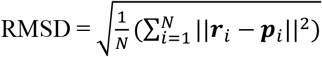

The image-to-atom regression architecture is summarized in **Fig. 2**.

## III. Data generation and forward model

In this section, we describe the data synthesis protocol (**Fig. 3**).

**Fig. 3.**
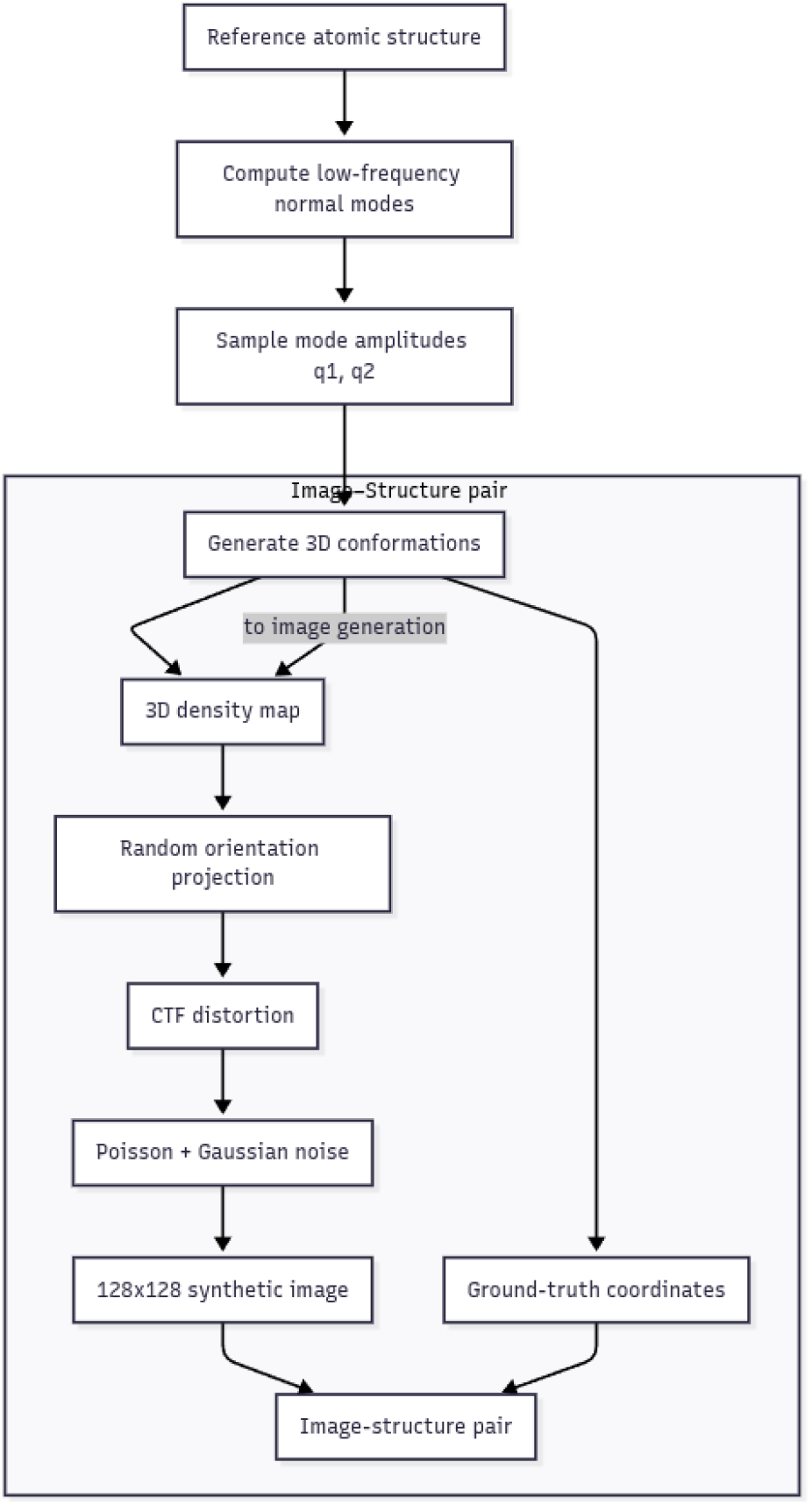
Synthetic data pipeline to produce pairs of synthetic images and atomic models, for testing the performance of the neural approach for predicting atomic models from cryo-EM images.

### A. Normal-Mode Conformational Sampling

We generate continuous conformational variability using low-frequency normal modes, which represent large-scale collective motions of a molecule around a reference structure [13]. Starting from a reference structure with *N* atoms at positions ***r***_*init*_, we compute normal modes ***a***_*i*_ using the so-called elastic network model [13, 14], and then, obtain elastically deformed structures by a linear combination of two low-frequency modes:

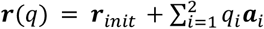

Amplitudes *q*_*1*_ and *q*_*2*_ are sampled from a random distribution. In this work, the choice of the two low-frequency normal modes is arbitrary. For adenylate kinase, we use normal modes 7 and 8 with the range of amplitudes [-120, 120] (note that the first six normal modes are never used as being related to rigid-body motions). For nucleosome core particle, we use normal modes 7 and 9 with the range of amplitudes [-150, 150]. These ranges produce smooth, continuous conformational variation while preserving structural integrity. For the reference structures, we use the Protein Data Bank (PDB) entries PDB:4AKE (adenylate kinase, chain A) and PDB:1KX5 (nucleosome core particle).

For both, adenylate kinase and nucleosome, 20,000 synthetic atomic models were generated and split into 15,000 training, 3,000 validation, and 2,000 test samples. Note that the obtained synthetic atomic models are in the same orientation and position as the reference structure used for their calculation. These rigid-body aligned conformations are then projected on randomly oriented image planes as explained next.

### B. Cryo-EM Image Simulation

A forward model converts each conformation into a synthetic single-particle cryo-EM image with a randomly sampled orientation, CTF modulation (defocus of 0.5 μm), and noise (SNR=0.1), following common cryo-EM simulation assumptions [1, 7, 9].

Images are generated at 128×128 pixels, with pixel sizes of 0.7 Å (adenylate kinase) and 1.3 Å (nucleosome). For both adenylate kinase and nucleosome, 20,000 images were generated and split into 15,000 training, 3,000 validation, and 2,000 test samples. The training, validation, and test datasets consist of the synthetic images and the corresponding synthetic atomic models.

## IV. Experiments

In this section, we describe experiments and results obtained with adenylate kinase and nucleosome (**Figs. 4-5**).

**Fig. 4.**
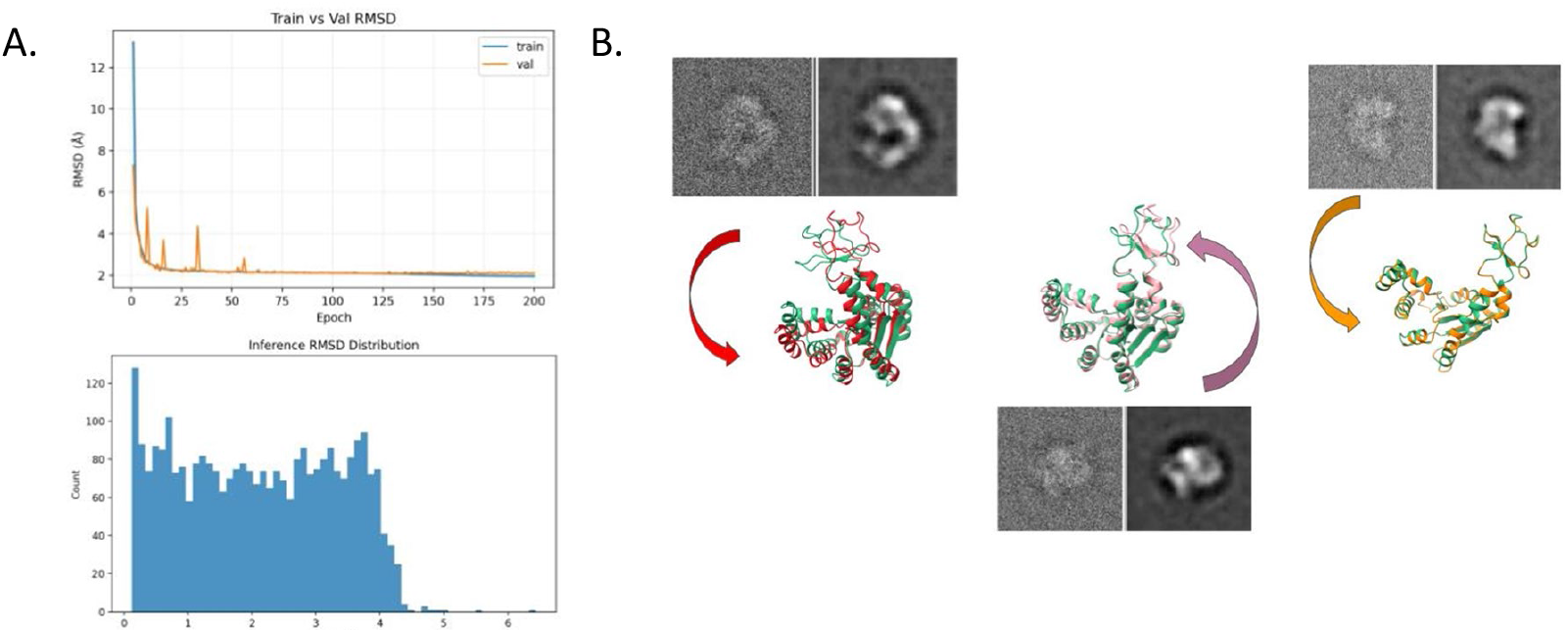
Adenylate kinase test case. A. Training and validation RMSD curves demonstrating stable convergence and absence of overfitting. B. Three examples of the inferred vs. ground-truth (green) structures (RMSD of 3.9Å, 2Å, and 0.2Å, for the red, pink, and orange reconstructions respectively). The corresponding images are also shown (ground-truth on the left and the prediction on the right in B).

### A. Training

Both subnetworks use cosine-annealed learning rate starting at 0.001. Gradient clipping at max norm 1.0 improves stability. Early stopping with patience 20-30 is used based on validation errors. Training is performed on an NVIDIA Quadro RTX 5000 (16 GB); typical runtime ranges from 3 to 12 hours, depending on whether the model is full-atom or coarse-grained, and depending on the molecular size.

### B. Inference

We evaluate atomic-model reconstruction accuracy using the RMSD (which is also used as a loss function for the second network) on 2000 images that were not used during training or test between the predicted and ground-truth atomic coordinates.

### C. Adenylate Kinase

On 2,000 test images, the image-to-atoms regressor achieves a mean RMSD of 2.11 Å, with a standard deviation of 1.22 Å, for the adenylate kinase’s all-atom representation (1,656 atoms). The RMSD distribution is concentrated, with 95% of test cases falling below 3.96 Å, and only two samples exceeding 5 Å. These results indicate that the learned image latents encode both pose and conformation with sufficient fidelity for atomic-level shape reconstruction. Training dynamics, inference reconstructions, and error distributions are shown in **Fig. 4**.

### D. Nucleosome Core Particle

For the nucleosome core particle’s coarse-grained representation (1,041 Cα and P atoms), the method achieves a mean RMSD of 0.80 Å with a standard deviation of 0.1 Å. Errors range from 0.2 to 2.3 Å, demonstrating that the approach scales to larger complexes and coarse-grained representations while maintaining near-atomic accuracy. The corresponding learning curves and RMSD distributions are shown in **Fig. 5**.

**Fig. 5.**
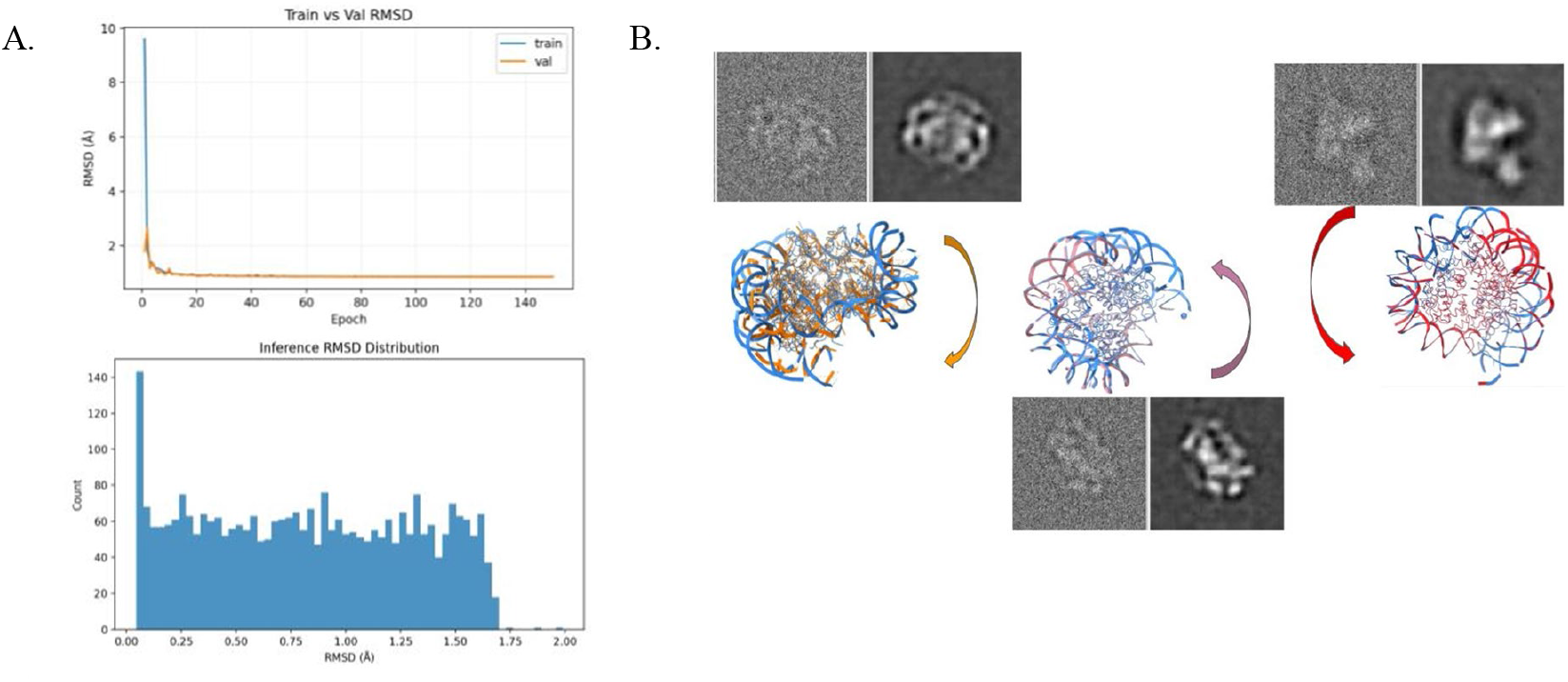
Nucleosome test case. A. Training and validation RMSD curves demonstrating stable convergence and absence of overfitting. B. Three examples of the inferred vs. ground-truth (blue) structure (RMSD of 2Å, 1.2Å, and 0.3Å, for the orange, pink, and red reconstructions respectively). The corresponding images are also shown (ground-truth on the left and the prediction on the right in B).

## V. Discussion

The results demonstrate that compact image latents extracted by a convolutional autoencoder can preserve sufficient information about both pose and conformation to permit subsequent regression of atomic coordinates. The image-to-atoms regression network effectively exploits these latents to recover structures without requiring pose estimation.

This approach is complementary to physics-based flexible-fitting approaches that compute one structural model per particle image based on costly MD simulations, such as MDSPACE [7], and could be combined for speed up of the analysis of large experimental cryo-EM datasets.

The study presented here was performed in a fully controlled, synthetic environment (the forward model is simplified and the ground-truth conformations are known and aligned) to evaluate the performance of the new method, before extending the study to experimental data. The RMSD between the predicted and ground-truth atomic coordinates achieved for both systems indicate that the mapping from image space to atomic coordinate space is learnable and that the mapping can capture subtle conformational variability.

In additional experiments (not shown here for space limit reasons), we have also considered mapping from image latents to atomic model latents, instead of the direct mapping from image latents to atomic coordinates. In this alternative approach (latent-to-latent mapping), atomic-model latents are obtained using an atomic-model autoencoder, consistent with supervised autoencoder formulations [15]. On the two test synthetic datasets, this alternative approach also achieved the mean RMSD of around 2 Å. These observations provide a proof-of-principle for coupling image and atomic-coordinate manifolds as a foundation for more advanced cryo-EM data analysis frameworks.

## VI. Conclusion and outlook

We presented a supervised framework for mapping noisy cryo-EM single-particle images to atomic coordinates by coupling image and structure manifolds. On synthetic datasets of adenylate kinase and nucleosome with conformations randomly oriented in 2D images but aligned in 3D structure space, the method achieves near-atomic accuracy of atomic-model reconstruction from images without orientation recovery. In future work, we will address the case of learning to predict conformations that are unaligned in 3D structure space, as they are unaligned in 2D images, which will allow pose recovery because the poses will be embedded in the atomic coordinates. Future work will also include combining this new approach with MDSPACE to enable fast determination of atomic conformational landscapes from large sets of experimental cryo-EM images.

## Acknowledgements

Acknowledgement

We acknowledge the support of the ANR (ANR-25-CE11-5519-01 to SJ), access to HPC resources of CINES and IDRIS granted by GENCI (AD010710998R3, AD010714089R1 to SJ), and the European Commission MSCA COFUND SOUND.AI and SCAI, Sorbonne University (PhD studentship to EA).

## Notes

### Competing Interest Statement

The authors have declared no competing interest.

